# The Tailtag System: Tracking Multiple Mice in a Complex Environment Over a Prolonged Period Using ArUco Markers

**DOI:** 10.1101/2024.11.07.622536

**Authors:** Vincent Coulombe, Arturo Marroquín Rivera, Sadegh Monfared, David-Alexandre Roussel, Quentin Leboulleux, Modesto R. Peralta, Benoit Gosselin, Benoit Labonté

**Affiliations:** Smart Biomedical Microsystems Labatory, Quebec City, Canada; CERVO Brain Research Centre, Quebec City, Canada

## Abstract

Despite recent advancements, safely and reliably tracking individual movements over extended periods, particularly within complex social groups, remains challenging. Traditional methods like colour coding, tagging, and RFID tracking, while effective, have notable practical limitations. State-of-the-art neural network-based trackers often struggle to maintain individual identities in large groups for more than a few seconds. Fiducial tags like ArUco codes present a potential solution by enabling accurate tracking and identity management, yet their topical application on mammals has proven difficult without frequent human intervention. In this study, we introduce the Tailtag system: a non-invasive, ergonomic tail ring embedded with an ArUco marker. This system includes a comprehensive parameter optimization guide along with practical guidelines on marker selection. Our Tailtag system demonstrated the ability to automatically and reliably track individual mice in social colonies of up to 20 individuals over a period of seven days without performance degradation, facilitating a detailed analysis of social dynamics in naturalized environments.

## 1 Introduction

The study of behavioral ethology necessitates the identification and tracking of individual movements within their environment over extended periods, requiring precise spatiotemporal information [1]. This becomes increasingly complex when applied to social dynamics, necessitating the monitoring of multiple members within a complex social group [2]. This complexity has limited our understanding of the different factors and conditions influencing social interactions and their dynamics in social groups [3].

A wide range of differentiation methods have been used in conventional laboratory setups, with pros and cons. Methods such as colour coding, ear-punching, and tattooing, while simple and cost-effective, have significant drawbacks. These methods can lead to mild issues, such as difficulties in reliable and automatic reading, as well as more severe concerns, including adverse physiological and behavioral consequences [4, 5]. As such, these methods can hardly be applied to the longitudinal monitoring of several animals together. Radio-frequency identification methods offer spatiotemporal tracking for large groups simultaneously [6]. However, their spatial resolution is limited by the density and positioning of antennas in the experimental environment. As a result, they are often incompatible with open-field setups and require carefully engineered layouts that force subjects into narrower spaces where identification can occur [3], potentially biasing behaviours. Additionally, RFID systems are cost-prohibitive for large-scale studies [7]. Recent advancement in deep learning allowed the development of effective solutions for tracking subjects with frame-level granularity [8, 9]. Unfortunately, even the current state-of-the-art in laboratory multi-subject tracking struggle to maintain accurate individual trajectories within large groups for more than a few seconds [10]. Tracking nearly identical subjects makes it difficult to recover tracking errors which end up accumulating over time, limiting the practical use of these solutions in laboratory settings [11]. Thus, reliably maintaining and recovering individual trajectories over extended periods, particularly within complex social groups and environments, remains an open challenge.

The mislabelling, identity swap and error propagation issues often affecting the tagless approaches described above can be mitigated by fiducial tags. Fiducial tags are objects placed in the field of view of an imaging system as a point of reference. Over the past years, several options have been developed, including ARTag [12] and ArUco [13]. ArUco tags are 2D binary-encoded fiducial patterns. Their design allows rapid, low latency detection (6D: 3D location and orientation) and identity of hundreds of unique tags. Behavioural monitoring using ArUco tags have been reported in ants, bumblebees, birds and cats [14, 15], providing important insights into the factors and conditions affecting social interactions and dynamics within social colonies. In these specific examples, ArUco tags were either printed and glued on the insects [15], combined to a backpack in bird studies [14] or on collars in cat studies [16]. Small devices designed to be mounted on top of head implants have also been recently developed to monitor engagement during cognitive tasks in mice [17]. However, applying ArUco tags directly on the mouse body to gather spatiotemporal information and infer social interactions within mouse colonies over prolonged periods of time has represented a challenge.

In this study, we developed the Tailtag system, a new approach for tracking individual mice in complex social groups within naturalized environments. The Tailtag is a 3D-printed black and white ArUco code imprinted on an 11 mm^2^ platform, easily positioned on the mouse’s tail and lasting for at least a week. To the best of our knowledge, the Tailtag system is the first solution to demonstrate, by itself, reliable tracking and re-identification capabilities in a large group (n=20) of mice over a prolonged period (7 days) in a challenging, enriched environment. We also utilized the system to analyze the evolution of social dynamics in our 20-mouse colony during the 7-day reliability trial. Our empirical results suggest that the Tailtag could be applied to conventional experimental setups or leveraged in conjunction with existing deep learning tracking solutions to improve the efficacy of long-term mouse social monitoring.

### 1.1 Materials and Methods

#### Animals

All animal experiments were carried out in male (n=20) C57BL mice (Charles River Laboratories, Kingston, NY) aged from 6 to 10 weeks old. All animal procedures followed the Canadian Guide for the Care and Use of Laboratory Animals and were approved by the Animal Protection Committee of Université Laval. Mice were housed in an enriched environment under a reverse 12-hour light/dark cycle starting at 7:00 AM, with food and water provided ad libitum.

#### Enriched Environment

We developed an ethological vivarium in which mice could live without human interference in large social groups. In total, 20 mice were housed simultaneously in this environment. The vivarium consists of a 1m x 1m x 0.5m open area made of ¾-inch thick plexiglass, providing 10,000 cm^2^ of living area. The vivarium is enriched with saw dust, wheels (n=2), tubes (n=4), wooden cubes (n=4) and ample Enviro-Dri nesting material (Figure 1a). Food and water is provided from food dispensers and water bottles. Two food/water sections are present in the vivarium (Figure 2f). The vivarium is in a quiet area with ambient noise below 50 decibels, it is maintained at 23 degrees Celsius with 60% humidity.

**Figure 1:**
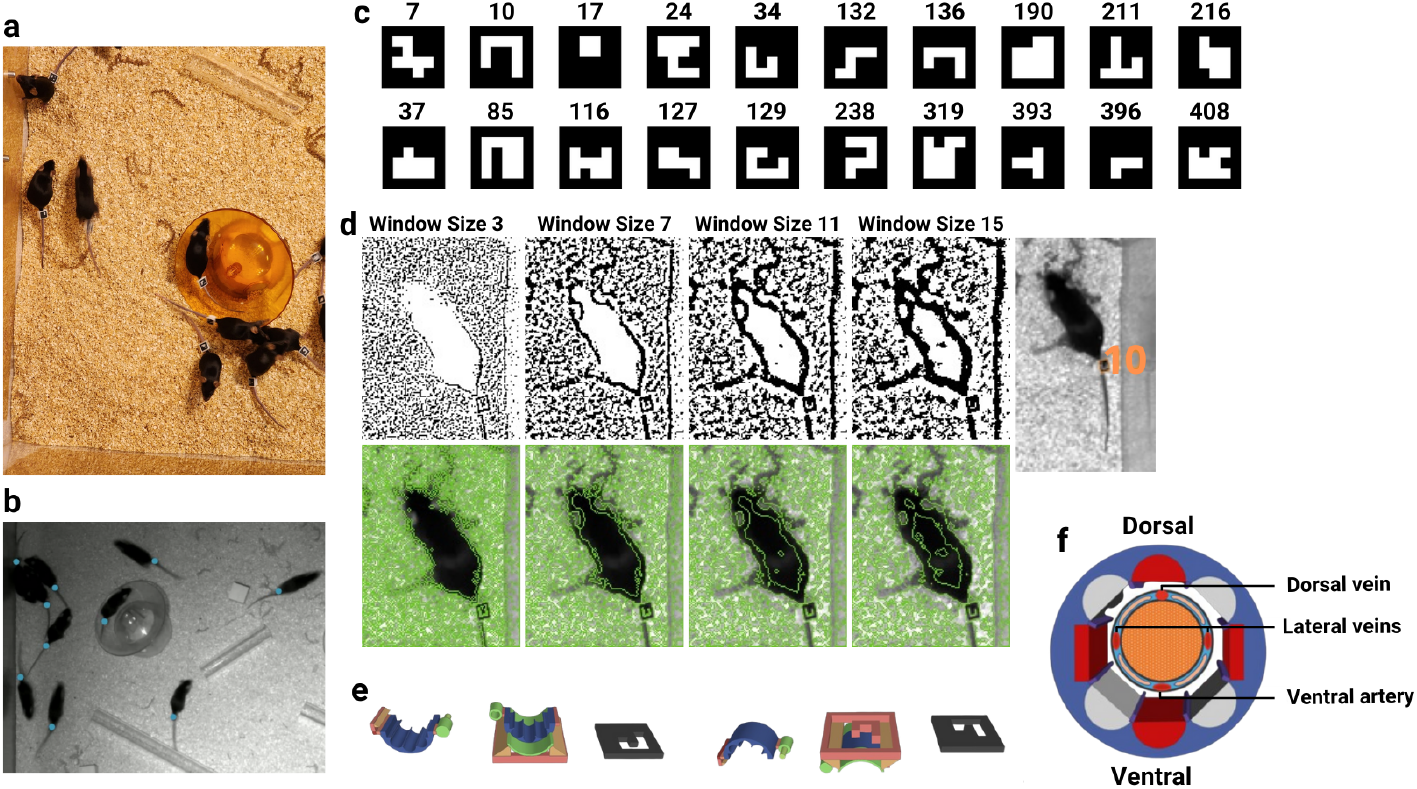
**a**, Close-up view of the enriched living area displaying the 600 mg, 11 mm^2^ Tail Tags in use. **b**, Tailtag’s coordinates inferred by our finetuned Yolox-pose algorithm. Akin to those used to automatically label the 5 040 000 tag’s locations leveraged during the reliability study. **c**, The selected Aruco markers and their binary maps. **d**, The thresholded image (top) and a visualization of the contours to filter (bottom), for different thresholding window sizes (in pixels) of 3, 7, 11, and 15. The thresholding window sizes impact the number of tags retained for the contour filtering (bottom) process. If the thresholding window sizes is too small, some markers’ border might break (top) causing it not to be detected. On the other hand, if the thresholding window sizes is too big, small markers might not get detected. **e**, The Tailtag’s 3D design. **f**, Schematic representation of a mouse tail with both dorsal and ventral arteries and lateral veins as well as the 3D design of the grooved cylinder to ensure normal blood circulation as monitor in our pilot study.

**Figure 2:**
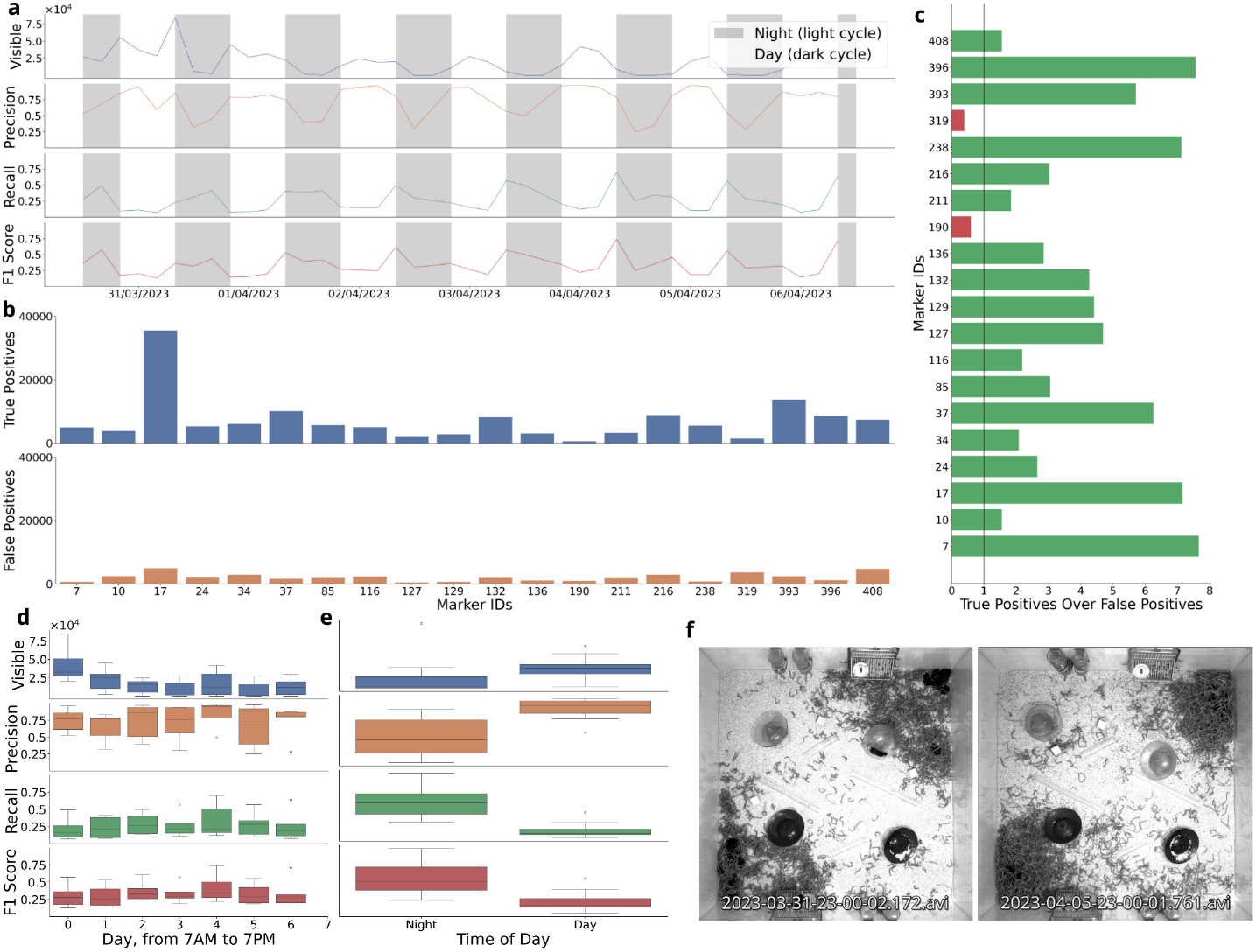
**a**, The number of visible tags, the recall, the precision and the F1 score evolution over the week-long trial. The nights, in grey, start at 7PM inclusively and end at 7AM exclusively. The count of visible tags in each recording ranged from about, 80000 at 7PM on the 2nd night to 2000 at 3 AM on the 6th night. **b**, The sum of all the true positives, or good detections, for each tag (top) and the sum of all the false positives, or bad detections, (bottom). **c**, the good positives to false positives ratios for each tag. A ratio inferior to one, highlighted in red, means that a marker was falsely detected more often than rightly detected. **d**, The number of visible tags, the recall, the precision and the F1 score distributions for each day of the week-long trial. **e**, The number of visible tags, the recall, the precision and the F1 score distributions during the day versus during the night. **f**, Top-down view of the vivarium for the 1st hour of the recording session (left) and on the 6th day of recording (right). We can qualitatively assess that the nests got considerably denser from the third day onward.

#### Image Acquisition

Mice were continuously video recorded over a 7-day period using an 8-megapixel monochrome Sony IMX265 camera with a 2064×1536 pixels resolution. The camera was positioned 215 cm above the vivarium and equipped with a 2/3 inch, 8mm, f/1.4 Manual Iris lens to capture the entire vivarium. This lens provided a wide field of view and a high aperture to ensure well-lit images even at fast shutter speeds. The vivarium was illuminated using a dual lighting scheme: infrared lights at night (light cycle) and LED lights during the day (dark cycle). To ensure the comfort of the animals, the LED lighting system was calibrated to maintain an illuminance level on the vivarium’s surfaces around 40 lux [18]. To minimize motion blur from the rapid movements of the mice, the camera was configured with a fast global shutter speed while maintaining good image quality. Given the moderate ambient light levels, a shutter speed of 1/1000 second was chosen.

The video camera was connected to a Dell laptop running licensed video recording software from Allied Vision. Video sequences were captured at 25 frames per second (FPS) with a resolution of 1536×1536 pixels. The recordings were automatically converted from .seq to .avi format on the laptop and segmented into 10-minute intervals to facilitate downstream analysis. These segments were then exported and stored on a DS1621+ Synology NAS, using a first-in, first-out (FIFO) system for time-scalable analysis. Specific segments, as part of the evaluation protocol, were automatically uploaded to a processing desktop for experimentation (computational resource usage). In total, the 7-day trial (168 1-hour recordings) generated 1.2 TB of uncompressed footage. All devices in the pipeline were connected via a TP-Link TL-SG108 network switch to maximize file transfer speeds.

#### Tailtags

We designed a 600mg portable 3D-printed device called the Tailtag (Figure 1e), which consists of two parts. The first part is a cylindrical component designed to attach the tag to the mouse’s tail. This cylinder measures 5mm in length and 10mm in width, with an inner diameter of 7mm. It features eight symmetric grooves and is divided into two hinged sections that can be clipped together around the mouse’s tail.

The dimensions and configuration of the Tailtag were determined based on a pilot study monitoring the growth of mouse tails in male mice (n=3) aged 6 to 8 weeks. At 6 weeks old, the average tail diameter was 3.43mm and increased to 3.87mm by 8 weeks. The 7mm inner diameter was chosen to minimize the risk of constriction and injury. The grooves inside the Tailtag are designed to accommodate the tail’s anatomy, which includes one ventral artery and three veins (two lateral and one dorsal; Figure 1f). These grooves ensure that blood circulation is not restricted, allowing for natural tail growth at various developmental stages. Bio-compatible silicone (Kwik-Sil, WPI Inc.) is used to fill the four diagonal grooves. This silicone adhesive keeps the Tailtag securely in place while allowing for natural tail growth.

The second part of the Tailtag is an ArUco marker, a 3D-printed, 11mm^2^ square platform on top of the cylinder. It features a black-bordered binary matrix that serves as a unique identifier within a predefined ArUco dictionary. For example, a 4×4 marker contains 16 bits for 216 unique black and white marker combinations. Both the markers and cylinders were printed using an LCD Anycubic Photon Mono X printer, which offers a 50-micron XY-pixel resolution by leveraging photopolymerization.

#### Tailtag Detection

Our detection system utilizes the OpenCV (V4.6.0.66) ArUco marker detection module [13]. The module identifies marker positions within video frames through a two-step algorithm. First, thousands of potential markers are detected using adaptive thresholding and a parameterized edge detection algorithm, which locates the markers by their borders. The potential markers are then filtered based on their contour’s shapes and sizes; those that do not approximate to a square shape are discarded. Second, the system identifies the IDs of the selected markers by analyzing their inner binary matrices (Figure 1c), by leveraging perspective transformations and Otsu thresholding to align and segment the black and white bits. These segmented bits are then examined to determine if the marker belongs to its respective dictionary. Both steps are highly configurable, and their respective parametrization processes are documented in the “*Selecting the System’s Parameters* “ subsection.

Detection performance, as defined by equations 1, 2 and 3, is significantly influenced by non-software-related factors, particularly the physical environment in which the tags are recorded. Factors such as inadequate lighting, tag obstruction, and rapid movement can impair tag visibility, leading to inaccurate or suboptimal trial classifications [19]. On the software side, the camera’s shutter speed must be optimized to reduce motion blur while preserving sufficient image quality. For more details, refer to the Image Acquisition subsection. Additionally, the parametrization of the ArUco detection system and the selection of its dictionary are critical. The subsequent subsections explain the system’s inner workings and our choices regarding acquisition parameters.

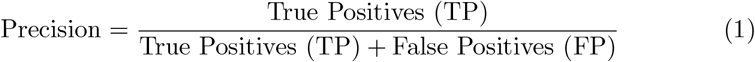

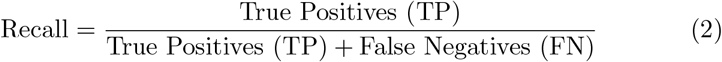

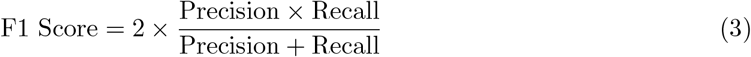

#### Tags Dictionary

The detection system provides several predefined tag dictionaries, varying from 4×4 to 7×7 bits, each containing between 50, 100, 250 or 1000 binary marker patterns. The tag size and the number of tags within the dictionary affect a crucial parameter: the inter-tag distance. This is the minimum Hamming distance between any two tags in the dictionary and impacts the system’s error detection and correction capabilities. A greater inter-tag distance results in more distinct tags, thereby reducing the potential for confusion.

Generally, smaller dictionary sizes and larger tag sizes increase the inter-tag distance. For instance, the minimum Hamming distance in a 50-tag dictionary with 4×4 bits is 4, whereas it decreases to 2 in a 1000-tag dictionary with the same bit size. Thus, if only 50 tags are required for an application, opting for the 50-tag dictionary is preferable, as it offers the maximum inter-tag distance and improves the system’s error resilience.

To thoroughly evaluate our detection system’s performance, we selected the predefined dictionary with the smallest Hamming distance between any two codes, specifically the 1000-tag 4×4 bits dictionary. Amongst these tags, we opted for a set of twenty tags listed in Figure 1c.

#### Selecting the System’s Parameters

As previously mentioned, the detection steps in our system are highly configurable. This customization enables users to fine-tune the algorithm for optimal performance. Most parameters involve a trade-off between detection performance and latency, as enhancement in detection accuracy often increases computational load. For long-term tracking, we selected a set of parameters that balanced robust performance with near real-time processing on our hardware. This approach prevents extensive computation times, making the system suitable for monitoring activities over days or even weeks. Refer to the Computational Resource Usage subsection for more details.

The baseline metric values for our parameter optimization experiments (Table 1) were obtained using the default parameters proposed by [13]. Table 1 presents the cumulative metric values achieved by incrementally tuning additional parameters. This tuning process involved a combination of logical reasoning, calculations, and systematic grid search, as documented in the following subsections.

**Table 1:** Cumulative parameter selection performances on our test recording.

| Modified Parameters (Cumulative) | Value | Precision (%) | Recall (%) | F1 Score (%) |
| --- | --- | --- | --- | --- |
|  |  | 90.0 | 11.0 | 19.6 |
| AdaptiveThreshWinSizeMin | 7 | **93.5** | 11.5 | 20.5 |
| AdaptiveThreshWinSizeMax | 23 |  |  |  |
| AdaptiveThreshWinSizeStep | 10 |  |  |  |
| MinMarkerPerimeterRate | 0.025 | 92.1 | 20.3 | 33.3 |
| MaxMarkerPerimeterRate | 0.035 |  |  |  |
| PerspectiveRemovePixelPerCell | 7 | 92.3 | 21.7 | 35.1 |
| PerspectiveRemoveIgnoredMarginPerCell | 0.35 | 92.0 | 23.8 | 37.8 |
| MinCornerDistanceRate | 0.15 | 92.1 | **24.0** | **38.1** |

##### Adaptive Thresholding

The first step in the marker detection algorithm, designed to detect potential markers by locating squared black borders within an image. This process experiment with gradually increasing thresholding window size (in pixels) to account for variations in lighting conditions, border sizes, and noise levels (Figure 1d). By default, the adaptive thresholding begins with a minimum window size (”*adaptiveThreshWinSizeMin*”) of 2 pixels, incrementally increasing by a step size defined by ”*adaptiveThreshWinSizeStep*” of 10 pixels, until it reaches the maximum window size (”*adaptiveThreshWinSizeMax* ”) of 23 pixels.

Qualitative assessment of our footage, as referenced in Figure 1d, indicates that an ”*adaptiveThreshWinSizeMin*” below 7 pixels does not improve detection, as most borders are barely visible or broken. This conclusion is quantitatively supported by an increase in every detection metrics when changing ”*adaptiveThreshWinSizeMin*” from 3 to 7 pixels (Table 1). However, lowering ”*adaptiveThreshWinSizeStep*” to 4 for allowing a threshold with window sizes of 11 and 15 pixels decreased detection metric values, so this parameter was left at its default value of 10 pixels. Similarly, adding a window size of 27 pixels unintuitively lowered metric values, so this option was also ruled out and the ”*adaptiveThreshWinSizeMax* ” was left at its default value.

##### Contour Filtering

The next step is to eliminate the non-marker candidates detected by the adaptive thresholding. Here, we aim to balance detection metrics and latency, as it is crucial to discard invalid candidates to save computational resources while avoiding overly strict filtering conditions that might discard real markers.

The first two tuned parameters, ”*minMarkerPerimeterRate*” and ”*maxMarkerPerimeterRate*”, determine the minimum and maximum size of a marker based on its perimeter relative to the maximum dimension of the input image. In our case, the average marker size captured by the camera is approximately 40×40 pixels. With a camera resolution of 1536×1536 pixels, the approximate perimeter rate of our markers is 40/1536 = 0.026. By grid searching values around this result, we found that a ”*minMarkerPerimeterRate*” of 0.025 and a ”*maxMarkerPerimeterRate*” of 0.035 provided the best detection metrics on our benchmark recording (Table 1).

##### Bits Extraction

In the final step of the detection algorithm, each remaining candidate’s bits are analyzed to determine their IDs. Most parameters in this step were left at their default values, as our grid searches found that they did not significantly impact detection performance. However, we identified that the ”*PerspectiveRemovePixelPerCell* ”, ”*PerspectiveRemoveIgnored-MarginPerCell* ”, and ”*minCornerDistanceRate*” are correlated with detection performance, at least on our test recording, and were worth tuning.

”*PerspectiveRemovePixelPerCell* ” represents the number of pixels for each bit on the tag. Knowing that our tags are 40×40 pixels with 4×4 bits plus borders, we set ”*PerspectiveRemovePixelPerCell* ” to 7, up from the default value of 4, which improved our detection metrics (Table 1). ”*PerspectiveRemoveIgnoredMarginPerCell* ” defines the margin of pixels per bit that counts toward final classification. A strong margin of 0.35 provided the best metrics. ”*minCornerDistanceRate*” represents the minimum distance between any pair of corners from the same marker relative to the marker perimeter. Increasing it to 0.15 from the default value of 0.05 slightly increased precision.

While many parameter configurations were tested, only those that impacted performance are presented. We did not use a corner refinement algorithm, as we did not deem the benefits of knowing the markers’ poses worth the extra latency cost.

#### Evaluation Protocol

The detection performance of our tags was assessed by comparing them against data automatically labelled by a finetuned YOLOX-pose neural network algorithm [20, 21] using the mmpose framework [22]. A small fraction of our recording frames (1000 images) was manually labeled to finetune the model. This finetuned model was then used to infer the Tailtag’s coordinates (Figure 1b). This approach enabled us to evaluate our tags’ detection performance against millions of labelled coordinates spanning over multiple days. Doing so, we were able to assess the detection performance over time and conclude that it does not deteriorate within a week.

Given the large volume of total recorded data (18,144,000 frames), evaluating the detection quality on every recorded frame would have been computationally intensive (taking weeks per evaluation) and redundant, as the detection context (the vivarium layout, the tag’s conditions and the lighting conditions) does not significantly change within minutes Therefore, we opted to assess detection quality using samples of our continuous week-long dataset. To minimize sampling bias, we uniformly selected recordings every 4 hours throughout the week, plus an additional video used for parameter tuning. Each of the resulting 43 samples consists of 15,000 data points, or 10 minutes of recording at 25 FPS. This offers a representative view of the entire dataset. The parameter tuning video was excluded from the final reliability evaluation to minimize potential overfitting, which could have skewed the results. We analyzed the first 10 minutes of each of the following hours: 11 PM, 3 AM, 7 AM, 11 AM, 3 PM, and 7 PM. Each day started when the LEDs in the room were on between 7 AM and 7 PM. Video sequences included day and nighttime recordings. Overall, this provided a total of 420 minutes (about 7 hours) of testing footage, uniformly distributed over the week.

##### Evaluation Criteria

A true positive was defined as an ArUco detection (x, y, w, h) encompassing a visible tag’s location (x, y). A false positive was defined as a detection not encompassing any tag’s location. Using these definitions, we calculated the classic classification metrics used to assess our detection performance using the equations 1, 2 and 3:

#### Computational Resource Usage

Using a single core with two threads, the OpenCV’s (V4.6.0.66) Aruco marker detection algorithm, configured with the parameters described in the Parameters Selection section, took an average of 29.66 milliseconds to process one 1536×1536 resolution frame on an Intel i7-10700k CPU running at 3.0 GHz. The running pipeline requires approximately 0.9 GB of RAM, and every 10 minutes of detection recording consumes between 1.3 KB and 1.2 MB of disk space, depending on the number of detected instances.

#### Behavioural study

In conjunction to assessing the reliability of our system over the course of a week, we analyzed the mice’s behavioural features using the 43 uniformly sampled 10-minutes recordings. We extracted the position of each mouse at every frame, following the MOT format [23]. Predefined areas of interest, including nests, feeders, water bottles, and wheels, were identified for tracking purposes. At any given time, if the x, y coordinates of a mouse’s tag fell within one of these areas, the mouse was considered to be inside that area until the tag was detected outside of it.

##### Outlier Detection

First, we examined the presence of outliers in our dataset and assessed the distribution of detections based on mouse identity and inverted circadian phase [day (dark phase): 7:00 AM to 7:00 PM] (see Supplemental Information: Fig. 1 A). We defined an outlier as any value exceeding the third quartile (Q3) plus one and a half times the interquartile range (IQR) of the detections, criterion met by one mouse (see Supplemental Information: Fig. 1 B).

#### Mouse to Area Interactions

A mouse-area interaction occurs when a mouse’s tag is detected within a predefined area and lasts until it is detected outside. We analyzed both the absolute and relative frequencies (percentages) of these interactions in each area of the vivarium. To assess variations in the proportions of mouse-area interactions, we conducted a Chi-square test on the counts of interactions over the total number of detections. Further, to identify in which areas the probability of a mouse-area interaction increases, we implemented a mixed multiple logistic model in which the dependent variable was a dichotomous factor indicating the occurrence or the absence (1 and 0, respectively) of a mouse-area interaction. To increase robustness, we considered a proper interaction only if a mouse interacted with an area for at least two seconds in a row. To build the latter, we assumed that if a mouse was detected inside an area during a second, it interacted with the area that second. The main independent variable was a categorical variable representing the four predefined areas, with the category exhibiting the fewest interactions (water bottles) used as the reference. We also included circadian cycle (dark vs. light phase) as a covariate, along with interaction terms between each area type and circadian cycle. To account for potential correlations between measurements from the same animal, we nested the measurements within mouse and period, allowing the intercept to vary randomly.

#### Mouse to Mouse Interactions

A mouse-mouse interaction is defined by the simultaneous presence of two mice in the same area. To quantify and segment these interactions, we estimated the pairwise Pearson correlation coefficients, in terms of number of interactions, between all pairs of mice and tested whether these estimates were significantly different from zero (no interaction). Further, since the feeders and the nests saw significantly more traffic than the other areas (Table 3), they were labelled as social hubs. Also, considering that the tags are easier to read within the feeding areas than within the nest areas, and that the top feeder is the area where the most tag detections occurred, we focused our downstream analysis on the top feeder. For this hub, we identified pairs of mice that co-interacted during the same periods more frequently than expected by chance, assuming a hypergeometric distribution using the R package ”cooccur.” Finally, we determined which pairs were more often significantly interacting across all the time periods. To achieve this, we created an index, adding one for every period in which the pair co-interacted and subtracting one for every period in which the mice were significantly discordant (they interacted less than expected by chance). We estimated the 99th and 1st percentile and considered the pairs in the extremes (head and tail of the distribution) as pairs that respectively spent significantly more time together or avoid each other.

All the analyses were performed in the R and Rstudio environment. The packages utilized are listed in the supplemental information (Supplemental information: List 1). For all the inferential tests herein performed, *α* = 0.05. When multiple testing was performed, we corrected the *p − value* with Bonferroni’s method.

## 2 Results

In this section, we present the results of our week-long reliability trial including the evolution of our detection accuracy (Figure 2a), which remained consistent over the duration.

### Parameters Selection

The parameter search, detailed in the Materials and Methods section, was conducted using a 10-minute recording at 25 fps for a total of 15,000 frames. This dataset was excluded from the final week-long evaluation set (see Evaluation Protocol for more details). Table 1 displays the cumulative performance of the parameter selections on our test dataset. Each row in the table inherits the parameter modifications from the previous rows, so the incremental improvements reflect the combined effect of all parameters, not just the latest changes. Globally, our analyses show that the Tailtags on their own are an effective tracking solution at a second-level granularity.

### Detection Accuracy

The level of performance obtained on our parameter tuning recording (see Parameter Selection) was sustained over a week-long evaluation trial. The average detection accuracies over the full 5 040 000 labelled data points are presented in Table 2.

**Table 2:**
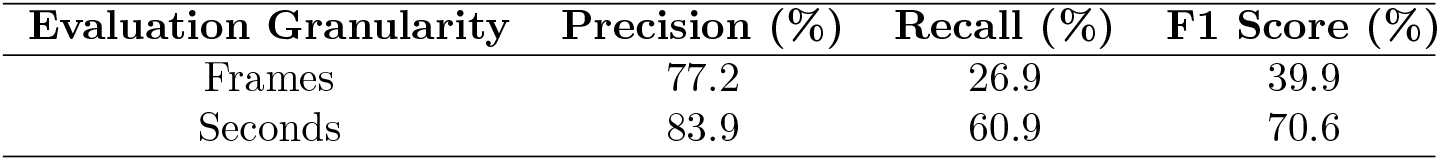
Detection accuracy over the week-long trial.

**Table 3:** Absolute and relative frequencies of mouse-area intersection.

| Area | Absolute Counts | % |
| --- | --- | --- |
| Top Nest | 47 477 | 25.0 |
| Bottom Nest | 40 045 | 21.0 |
| Top Feeder | 36 569 | 19.0 |
| Bottom Feeder | 18 739 | 9.9 |
| Top Right Wheel | 3 384 | 1.8 |
| Bottom Left Wheel | 2 663 | 1.4 |
| Top Left Wheel | 2 336 | 1.2 |
| Top Right Water Bottle | 1 421 | 0.7 |
| Bottom Right Water Bottle | 1 369 | 0.7 |
| Bottom Left Water Bottle | 906 | 0.5 |
| Top Left Water Bottle | 777 | 0.4 |
| Bottom Right Wheel | 464 | 0.2 |

Although the reported metrics are at frame-level granularity (Table 2, Frames), we recognize that pinpointing a mouse’s location with 1/25th of a second precision may be unnecessarily precise. In practice, users might be more interested in detection accuracy over a broader time span, such as one second. As mentioned in the Evaluation Protocol section, our recordings were captured at 25 frames per second (fps), meaning the algorithm had 25 opportunities to detect a given tag each second. Leveraging this, we estimated the accuracy metrics over one-second intervals (Table 2, Seconds). In this context, the recall metric improved significantly, as it now represents the percentage of visible mice detected at least once within each 25-frame segment of the week-long recording. This second-level recall better aligns with the qualitative performance users are likely to experience in practice.

### Individual Marker Precision

We tested whether the detections could be biased toward specific tags and whether this would affect accuracy. Figure 2b highlights notable differences in detection frequency for each tag in our complex environment. For example, tags 17, 37, and 393 were detected far more often than tags 190 and 319, suggesting that marker IDs could be yet another important parameter to optimize for detection frequency. However, true positive detection alone can be misleading, as some mice might spend more time hidden than others. To better assess detection quality, we analyzed the true positive to false positive ratio for each tag. Figure 2c shows that 75% (15/20) of the tags have a ratio above 2, 55% (11/20) above 3, and 20% (4/20) reach a ratio above 7. Only the two tags with the lowest detection rates (190 and 319) have ratios below 1, suggesting that marker ID selection likely impacts detection performance.

### Detection Over Time

To assess whether the system could provide reliable detections over a 7-day period, we analyzed 7 hours of footage, evenly distributed across 7 days (see Materials and Methods for details). Figure 2d) shows the daily aggregated results for our three primary metrics, along with the number of visible tags. To assess the consistency of performance metrics throughout the week, we conducted a one-way analysis of variance (ANOVA), using each day’s metrics as levels. The F1 score (one-way ANOVA, *p* = 0.53), recall (one-way ANOVA, *p* = 0.49), and precision (one-way ANOVA, *p* = 0.58) all adhered to the null hypothesis *H*_0_ (Equation 4), showing no statistically significant variation throughout the week. These findings suggest that the tag integrity is maintained over time, allowing for accurate and efficient detection for periods of at least 7 days.

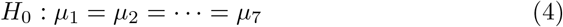

Where:

- *μ*: Level means.

Our analysis also revealed a significant decrease in the number of visible tags during the second day, dropping from approximately 60,000 on day 1 to an average of 17,000 from day 3 (one-way ANOVA, *p − value* = 0.004). A visual investigation of the video footage revealed that this drop is likely due to the mice starting to shape their environment by building denser, thus more occlusive nests (Figure 2f) which could have impacted the number of visible tags within those areas. From the third day onward, the number of visible tags per day does not vary significantly (one-way ANOVA, *p − value* = 0.18), indicating that a steady state was reached. This is supported by qualitative analysis, showing no significant changes in nest shape or density after the second day. Globally, although the number of mice in the vivarium did not change, fewer mice were visible from days 3, resulting in fewer detected tags.

### Day versus Night

We then tested the system’s capacity to detect tags during nighttime (light cycle). As depicted in Figure 2e, there is a notable reduction in the number of visible tags at night (10,000 on average at nighttime and about 26,000 at daytime), suggesting that fewer mice are roaming during this period of the day. This decrease inevitably leads to fewer inter-peer obstructions on average. Additionally, our qualitative analysis of nighttime footage reveals that mice active at night tend to sprint less, which reduces motion blur and further improves the accuracy of tag readings. Overall, our analysis suggests that while there is no significant difference between the metrics for night and day, the data is skewed towards a better recall at night and a better precision during the day. This suggests that both lighting and the mouse behaviour play a role in the system’s effectiveness.

### Behavioural Study

In conjunction to assessing our system reliability, we also assessed its ability to infer behavioural features within social groups in complex, naturalized environments. For this, we used our 43 uniformly sampled 10-minute recordings, yielding a total of 14,421,860 rows (frame per period per mouse), of which 232,742 contained at least one Tailtag detection.

Leveraging this positional dataset, we aimed to address ethological questions, such as determining zone preferences among the mice. Figure 3a illustrates the predefined zones along with each mouse’s activity level during both light (yellow) and dark (blue) phases, showing that activity was not confined to any specific zone. Figure 3b demonstrates that these zone preferences were dynamic, fluctuating not only with the circadian cycle but also throughout the study period. This suggests that preferences for specific areas varied over time, potentially influenced by evolving social dynamics. Additionally, certain zones, particularly the feeders and nests, showed significantly higher levels of activity (Table 3). Figure 3c reveals that social interactions in these predefined areas varied over time, displaying highly polarized co-occupancy levels in some zones. Notably, the feeders exhibited periods with either very few or many mice present simultaneously. This pattern indicates that feeding behaviour likely has a social aspect, positioning feeders as key social hubs. Based on these insights, we focused on the top feeder, the most frequented social hub, to investigate whether specific groups of mice formed for feeding and whether the composition of these groups remained stable over time.

**Figure 3:**
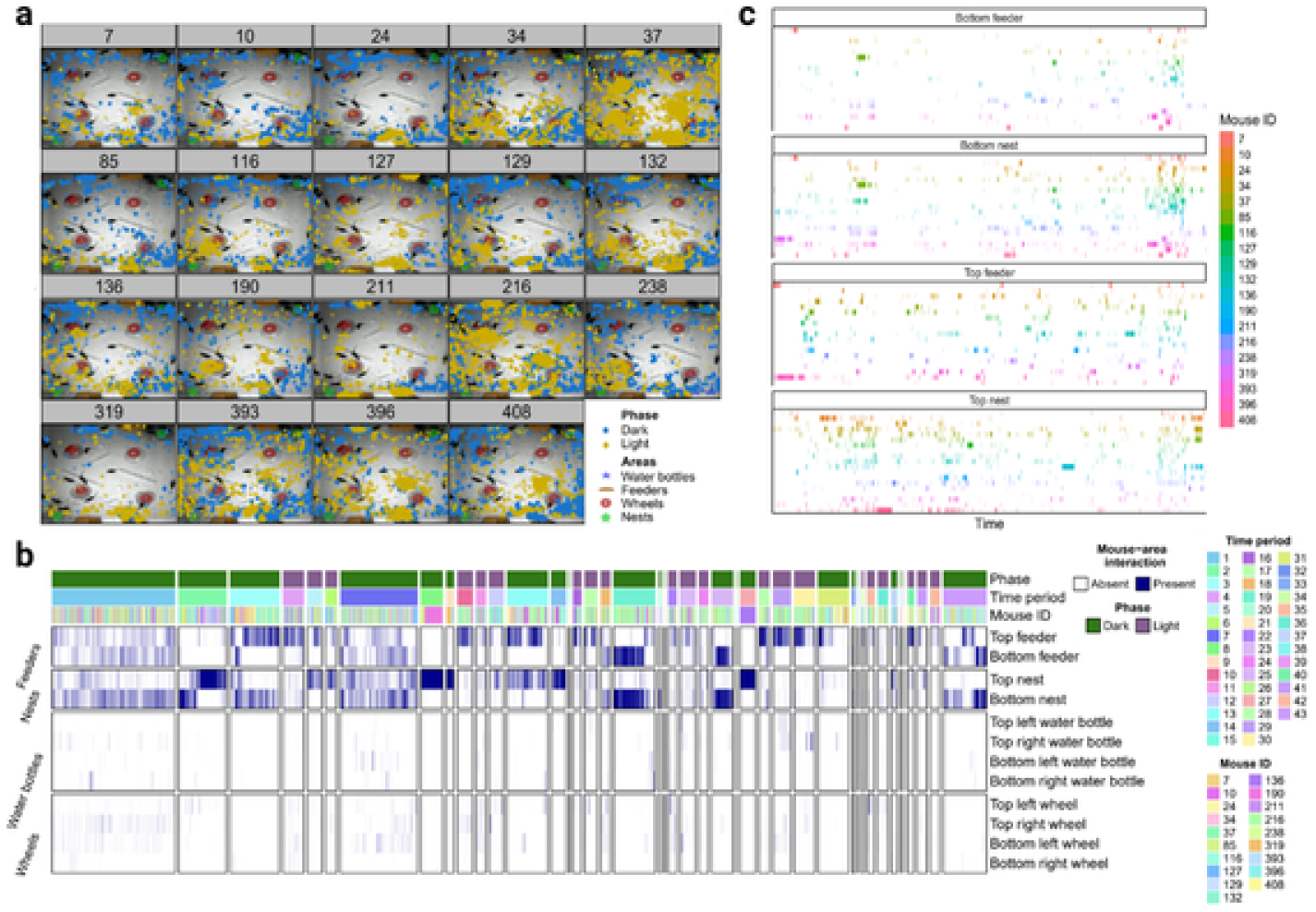
Evolution of the mouse-area interactions over time. **a**, Scatter plot showing the position of the mice for every frame during the 43 periods of 10 minutes. The plots are based on the coordinates of the individuals in the vivarium and were concatenated in chronological (date time) order. **b**, Heatmap illustrating mouse detection within each area over time. The total of detection per seconds within each of the four principal social hubs throughout the duration of the study. **c**, The 2-second interactions (see methods) shown for the four principal hub areas. The x-axis is time chronologically ordered.

Figure 5a illustrates overall social preferences in the vivarium, represented by pairwise correlations between mice based on cumulative co-detections of at least two seconds at the top feeder. For instance, mice 24 and 129 showed a significantly positive correlation at this feeder, while mice 211 and 216 exhibited no notable interactions with others. This analysis revealed five distinct social groups that interacted more frequently at the top feeder, consisting of 8, 5, 4, 1, and 1 member(s), respectively. While this analysis provides insight into general social interactions between pairs, further examination is needed to understand the temporal fluctuations and dynamics of these interactions. To achieve this, we analyzed the overall frame co-occurrence of the mice throughout the week specifically at the top feeder, depicted in Figure 4, which reveals variations in social dynamics over the recorded periods. Notably, the first period exhibited higher activity levels, as the nests had not yet been fully compacted by the subjects (Figure 2f).

**Figure 4:**
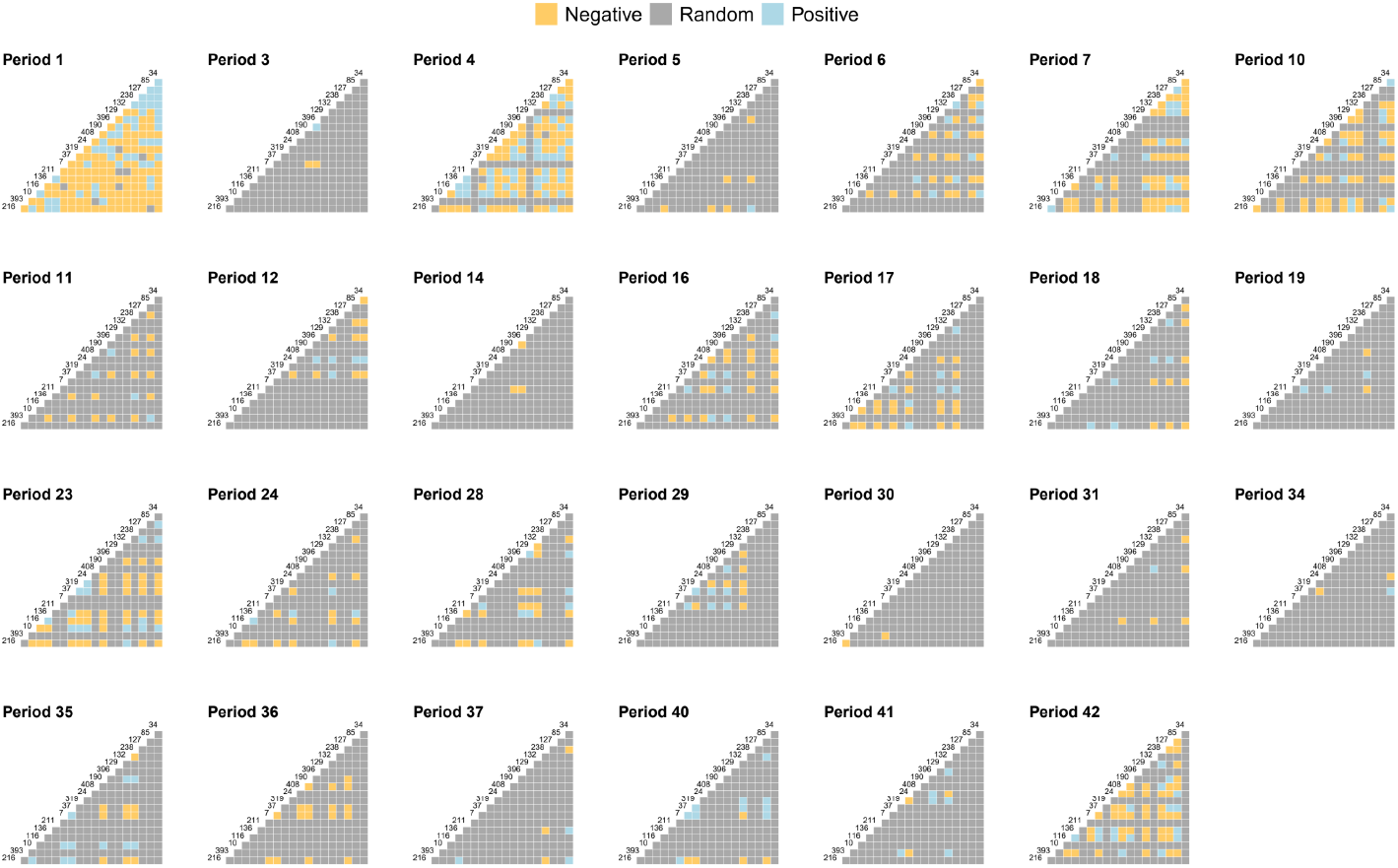
Co-occurrence matrices for all the well-powered periods of time. The co-occurrence was determined per frame.

**Figure 5:**
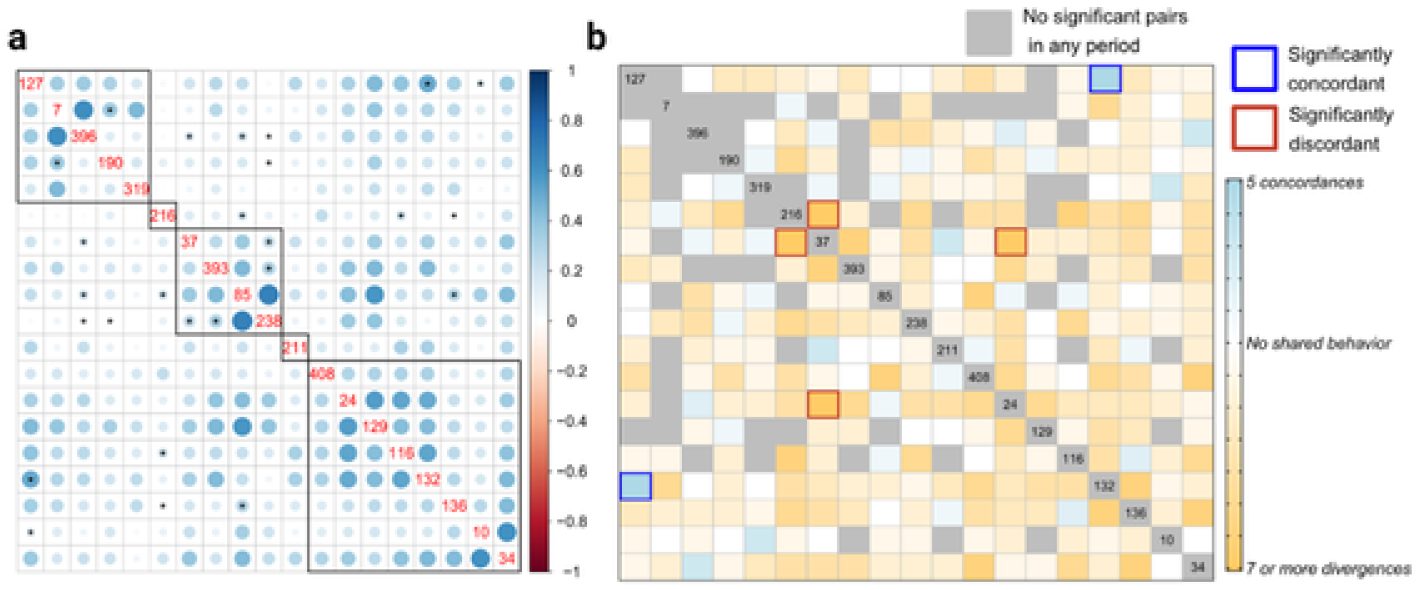
**a**, Correlation matrix of the number of mouse-area interactions based on the Pearson’s correlation coefficient. *Showing the Bonferroni adjusted *p − value*. Hierarchical clustering was performed to generate five groups, which are represented by the black squares **b.** Matrix of the index computed according to the number of periods in which a mouse significantly interacted or rejected another mouse at the top feeder. The red squares depict the pairs that spent more time or less time together at the top feeder throughout the study,

To further quantify these evolving social dynamics, we calculated an interaction index (see Mouse to Mouse Interactions in the Methods section for details) to assess whether pairs of mice were more likely to interact or avoid each other over the week. Supplemental Figure 2 presents the distribution of this index, distinguishing between pairs that showed a tendency to interact (head) and those that exhibited avoidance (tail). Using these indices, we determined whether the probability of interactions or avoidance among mice at the top feeder was higher than expected (Figure 5b). Notably, mice 132 and 127 showed a clear preference for interacting together, while mice 216 and 24 displayed avoidance towards mouse 37. Additionally, mouse 393 appeared to avoid specific partners and randomly engage with others, highlighting the complex and dynamic nature of social structures within the colony. This analysis supports the existence of social groups at the top feeder and suggests the possible exclusion of certain members. These findings confirm that social preferences within a colony are intricate and evolve over time.

## 3 Discussion

In this study, we evaluated the reliability of ArUco markers for tracking and reidentifying mice using the Tailtag system: a 3D-printed ArUco marker mounted on an 11 mm^2^ platform, ergonomically attached to the mouse’s tail. The system proved durable and effective, maintaining tracking efficacy across a large group of 20 mice over a 7-day period. Additionally, we used the Tailtag system to observe the evolution of social dynamics within the colony during the trial, offering insights into the system’s reliability within an enriched, socially complex environment.

Existing methods for tracking and distinguishing mice present several limitations. Colour coding and fiducial tagging require frequent maintenance [5], tattooing can pose health risks, and ear punching is difficult to monitor from a top-down, social environment view [4]. RFID systems, though widely used, lack the spatial precision needed for accurate tracking in open-field settings [3]. To address these gaps, we developed the Tailtag system: a non-invasive, ergonomic tail ring embedded with an ArUco marker. This system includes a comprehensive parameter optimization guide for the OpenCV ArUco module (v4.6.0.66) [13] along with practical guidelines on marker selection, hardware, lighting, and camera setup. The Tailtag system allows precise, continuous multi-mouse tracking in complex environments both day and night, sustaining effectiveness for at least a week without harming subjects or requiring frequent human intervention. This approach enables the automated collection of highly detailed spatiotemporal data with second-level granularity, supporting large-scale behavioural analysis.

Monitoring a large group of socially interacting subjects in a complex environment over an extended period with high spatiotemporal accuracy allows us to overcome some of the limitations faced by previous behavioural studies of smaller scope [1, 2]. This approach has enabled us to quantify the significance of specific social hubs and observe the evolution of social dynamics within them. Notably, we identified five distinct social groups. Our analysis also revealed interactions and avoidance patterns between specific pairs of mice within the most active social hub, the top feeder, highlighting examples such as the highly interactive pair of mice 132 and 127, as well as cases of selective avoidance, like mice 216 and 24 avoiding mouse 37. Overall, our findings indicate that while the zone preferences and peer associations among the mice change over time, certain groups and pairwise interactions consistently form within the social colony.

The Tailtag system offers significant benefits over existing methods for acquiring behavioural data automatically. Notably, since the performances do not degrade over at least a few days, a high percentage of the recorded footage end up being usable, contrasting with other available methods [3]. Also, the Tailtags offer high spatiotemporal accuracy even in wide open-fields. A limitation met systematically by RFID-based methods [3]. As such, researchers using RFID to collect behavioural data have to cleverly design specific narrow areas in the subjects’ environment where the detections can occur, giving the researchers datapoints only for those areas. Additionally, topical fiducial tags or colour codings methods, often leveraged in hairless animals [15], tend to require frequent reapplication in rodents [5]. In contrast, Tailtags are securely and ergonomically attached to mice tails, maintaining readability without frequent human intervention. While Tailtags have some challenges, such as needing exhaustive experimentation for both optimal parameter tuning and marker selection, our in-depth configuration guide as well as our global marker selection recommendations assists in streamlining these processes. Adequate lighting and a high resolution, fast global shutter camera are also recommended for optimal performance. Once properly configured, the Tailtag system provides a precise, hands-off solution for safely tracking multiple mice in complex, open environments.

Tracking and analyzing behaviour within large social groups is inherently challenging, especially when subjects are visually similar, as with mice. While past research has leveraged machine learning algorithms, these methods often struggle to balance accuracy and scale [8, 9]. This is because, vision-based animal tracking systems frequently suffer from identity switches when two mice cross paths and briefly occlude each other. Due to the subject’s visual similarity, these errors are often irrecoverable and degrade tracking accuracy over time. In contrast, human tracking systems leverages subject’s wearable accessories, such as clothes or bags, to visually differentiate individuals [24]. Drawing on this idea, we designed the Tailtag as a non-invasive, mouse wearable accessory, providing a reliable means of re-identification. This novel system has potential for integration with neural network trackers to mitigate their aforementioned re-identification issues,

### 3.1 Conclusion

In this study, we introduced the Tailtag system, a novel method for tracking individual mice in naturalized social groups. The Tailtag consists of a 3D-printed black and white ArUco code mounted on an 11 mm^2^ platform that attaches to the mouse’s tail. To our knowledge, this is the first physical re identification system to show, by itself, reliable mice tracking and re-identification within a large group (n=20) of interacting individuals over an extended period with second level spatiotemporal precision. Specifically, we showed no degradation in both precision and recall metrics over the course of a 7-day trial. Leveraging the collected data, we uncovered significant social dynamic shifts throughout the week and identified key social hubs and peer preferences within the enriched environment. Although the Tailtags are reliable, they lack the ability to provide detections at a frame-level granularity, since most tags require about 25 frames to be detected. This suggests that the Tailtag system could be leveraged by experiment needing second-level granularity. Alternatively, the Tailtags could be paired with neural network tracking solutions to enable long-term social monitoring of mice while still maintaining frame-level granularity, which would enable fine-grained behavioural monitoring with long-term reliably. Thus, giving the field an effective method for monitoring large and complex social groups with precise spatiotemporal information over a prolonged period.

