## Supplemental Material for "The Tailtag System: Tracking Multiple Mice in a Complex Environment Over a Prolonged Period Using ArUco Markers"

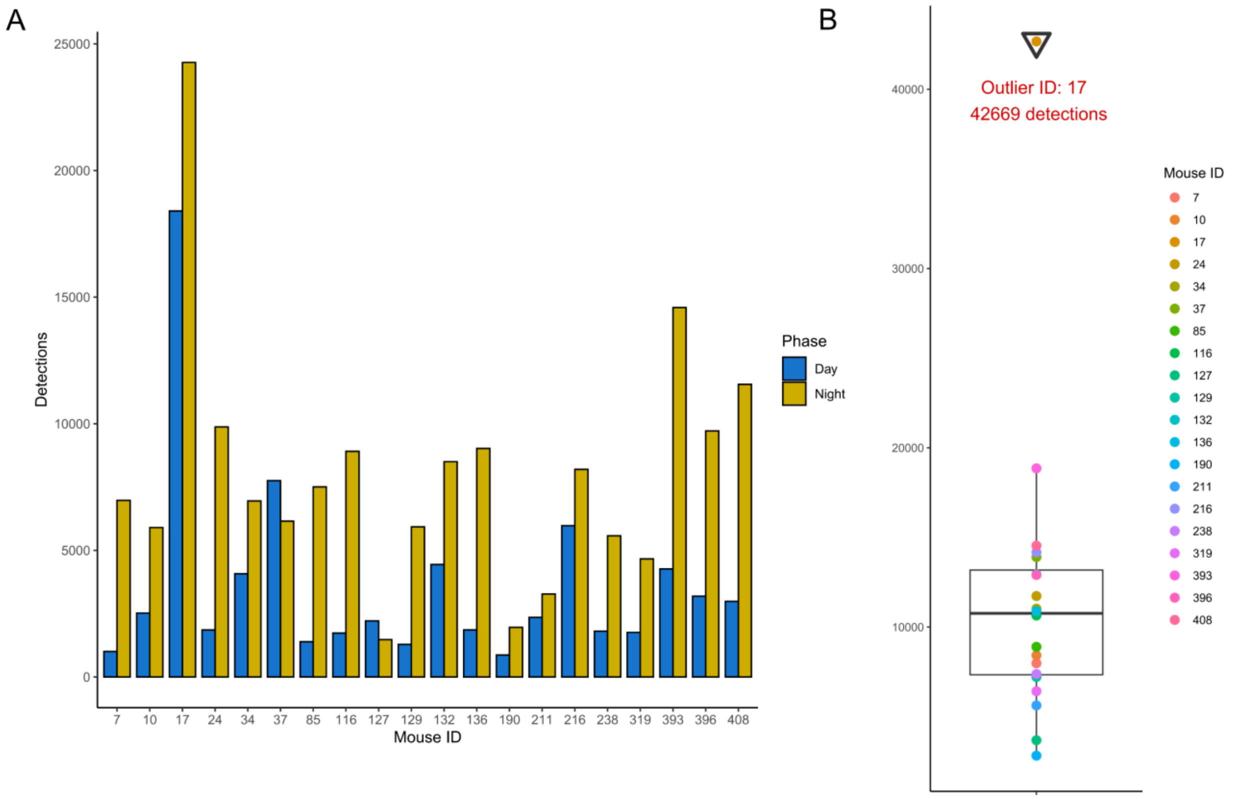


**Figure S1: A**. Barplot showing the number of detections for every mouse according to the circadian cycle. **B**. Boxplot showing the distribution of detection per mouse and the outlier according to the stablished criterion (see methods).

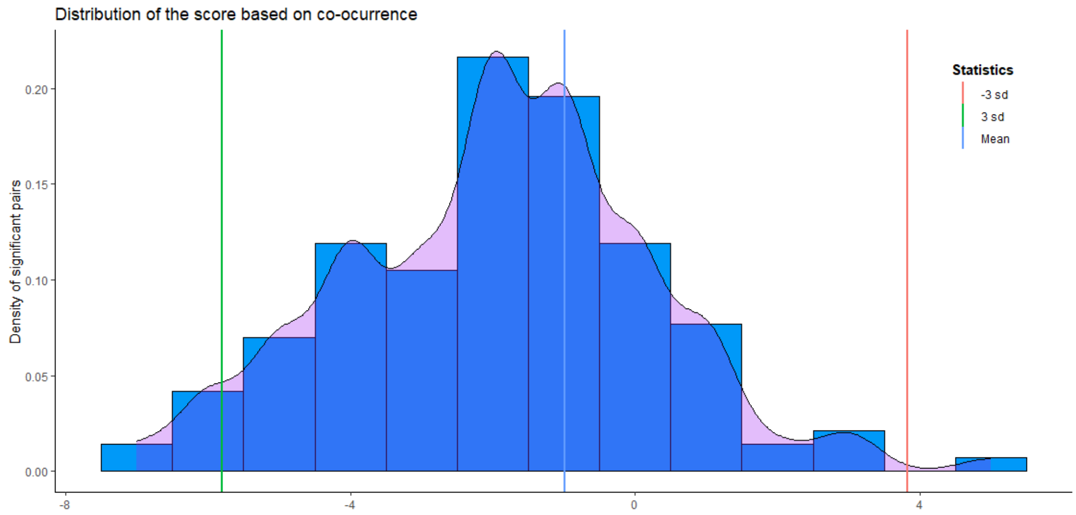


**Figure S2:** Distribution of the social index built in order to evaluate mouse-mouse interactions and avoidance (see methods).

**List 1:** R packages used in the study

- ggplot2
- dplyr
- gtsummary
- tidyr
- stringr
- ComplexHeatmap
- data.table
- sjPlot
- corrplot
- circlize
- cooccur
- ggpubr
- forcats
- lme4
- lmerTest

| | Mouse-object interaction per second | | |
|---|---|---|---|
| *Predictors* | *OR* | *CI* | *Adjusted – p value* |
| (Intercept) | 0 | 0.00 – 0.01 | **<0.001** |
| Feeders | 22.23 | 19.63 – 25.17 | **<0.001** |
| Nests | 55.72 | 49.36 – 62.88 | **<0.001** |
| Wheels | 2.97 | 2.59 – 3.41 | **<0.001** |
| Circadian cycle (Dark phase) | 0.81 | 0.66 – 0.99 | 0.32 |
| Feeders x Dark phase | 0.83 | 0.71 – 0.99 | 0.26 |
| Nests x Dark phase | 0.23 | 0.19 – 0.27 | **<0.001** |
| Wheels x Dark phase | 1.03 | 0.86 – 1.24 | 0.72 |
| Marginal $R^2$ / Conditional $R^2$ | 0.318 / 0.404 | | |

**Table1:** Multiple mixed logistic model where the occurrence of a 2 second interaction is the dependent variable and the main independent variable is a categorical variable including the 4 object type with the "Water bottles" as the reference. It also includes circadian phase (Dark vs Light phases) as covariates and interaction terms between the objects and the circadian phase. Measurements are nested the within mouse and period allowing the intercept to be random. The p value is adjusted by Bonferroni's method considering all each test performed in the model.
